# A paradigm for ethanol consumption in head-fixed mice during prefrontal cortical two-photon calcium imaging

**DOI:** 10.1101/2023.07.20.549846

**Authors:** Anagha Kalelkar, Grayson Sipe, Ana Raquel Castro E Costa, Ilka M. Lorenzo, My Nguyen, Ivan Linares-Garcia, Elena Vazey, Rafiq Huda

**Affiliations:** WM Keck Center for Collaborative Neuroscience, Department of Cell Biology and Neuroscience, Rutgers University – New Brunswick, 604 Allison Road, Piscataway NJ, 08904 USA; Department of Brain and Cognitive Science, Picower Institute for Learning and Memory, Massachusetts Institute of Technology, 43 Vassar Street, Cambridge MA, 02139 USA; Department of Biology, The University of Massachusetts Amherst, 611 North Pleasant Street, Amherst MA, 01003 USA

**Keywords:** Alcohol, prefrontal cortex, anterior cingulate cortex, ethanol, head-fixed, binge drinking, two-photon, calcium imaging

## Abstract

The prefrontal cortex (PFC) is a hub for higher-level cognitive behaviors and is a key target for neuroadaptations in alcohol use disorders. Preclinical models of ethanol consumption are instrumental for understanding how acute and repeated drinking affects PFC structure and function. Recent advances in genetically encoded sensors of neuronal activity and neuromodulator release combined with functional microscopy (multiphoton and one-photon widefield imaging) allow multimodal *in-vivo* PFC recordings at subcellular and cellular scales. While these methods could enable a deeper understanding of the relationship between alcohol and PFC function/dysfunction, they require animals to be head-fixed. Here, we present a method in mice for binge-like ethanol consumption during head-fixation. Male and female mice were first acclimated to ethanol by providing home cage access to 20% ethanol (v/v) for 4 or 8 days. After home cage drinking, mice consumed ethanol from a lick spout during head-fixation. We used two-photon calcium imaging during the head-fixed drinking paradigm to record from a large population of PFC neurons (>1000) to explore how acute ethanol affects their activity. Drinking modulated activity rates in a subset of neurons on slow (minutes) and fast (seconds) time scales but the majority of neurons were unaffected. Moreover, ethanol intake did not significantly affect network level interactions in the PFC as assessed through inter-neuronal pairwise correlations. By establishing a method for binge-like drinking in head-fixed mice, we lay the groundwork for leveraging advanced microscopy technologies to study alcohol-induced neuroadaptations in PFC and other brain circuits.

**Highlights:**

- C57BL/6J mice voluntarily consume ethanol to binge-like levels during head-fixation, with females consuming more ethanol than males.
- Mice show differences in frontloading and licking behavior for head-fixed ethanol and sucrose consumption.
- Head-fixed paradigm enables two-photon calcium imaging in the anterior cingulate cortex subdivision of the prefrontal cortex.
- Acute ethanol increases and decreases single neuron activity at fast (seconds) and slow (minutes) time scales but does not alter pairwise correlations between neurons.

## 1. Introduction

Binge drinking is defined by the National Institute on Alcohol Abuse and Alcoholism as a pattern of alcohol consumption leading to blood ethanol concentrations (BEC) of >80mg/dL. Binge drinking poses significant health risks including severe potential acute effects such as bodily injury, automobile accidents, and death from overdose. Repeated episodes of binge drinking also lead to adverse consequences affecting the heart, liver, and other organ systems. Moreover, binge drinking is a major risk factor for developing alcohol use disorder (AUD), a chronically relapsing condition characterized by compulsive alcohol seeking and consumption. Neural mechanisms that promote binge drinking and the neuroadaptations elicited by repeated alcohol intake remain incompletely understood but are important to resolve in order to identify novel therapeutic targets for curbing excessive drinking and the associated health burdens.

The prefrontal cortex (PFC) is an associative region of the brain with canonical roles in executive function and cognition, including decision making, attention, reward/punishment processing, social behavior, and other functions (Alexander and Brown, 2011). Consistent with the association of AUD with dysfunction of these higher-order processes (Bechara et al., 2001; George and Koob, 2010; Ridderinkhof et al., 2002), the PFC is highly implicated in AUD in humans (Goldstein and Volkow, 2011; Wilcox et al., 2014) as well as in non-human primate (Jedema et al., 2011) and rodent models (Halladay et al., 2020; Pava and Woodward, 2014; Robinson et al., 2019; Salling et al., 2018). Significant evidence describes mechanistic changes in PFC structure and function following acute or chronic alcohol exposure (Cannady et al., 2021; Lu and Richardson, 2014), with changes typically assessed in separate cohorts of animals at various time points throughout the alcohol exposure paradigm. However, little is known about the effects of alcohol on the real time neurophysiology of individual neurons and neuronal networks in intact circuits of behaving animals voluntarily consuming alcohol. Moreover, understanding how neuronal processing in PFC networks on the sub-second time scale relates to voluntary drinking, and tracking this relationship across days and weeks of drinking, is crucial for identifying the key neuroadaptations underlying the development of AUD.

Research in preclinical animal models, wherein animals voluntarily consume ethanol to binge-like levels, has generated significant mechanistic information on the bidirectional relationship between neuronal activity and ethanol consumption (Holleran and Winder, 2017; Siciliano et al., 2015). Several paradigms have been used successfully for achieving binge-like BECs in animal models, including procedures for home cage drinking in mice (Crabbe et al., 2011; Rhodes et al., 2005). Though these settings represent more naturalistic contexts, they limit the technical approaches that can be used to measure and manipulate *in vivo* brain dynamics. By leveraging emerging tools for activity recordings and manipulations in freely behaving mice (for e.g., electrophysiology (Cannady et al., 2020; Linsenbardt and Lapish, 2015), fiber photometry (Gioia and Woodward, 2021; Liu et al., 2020) and chemogenetics (Dao et al., 2021; Giacometti et al., 2020; Rinker et al., 2017)), recent studies have identified how the molecular, cellular, and circuit properties of neurons and glia in defined brain circuits for reward, motivation, and stress both affect and are affected by binge-like consumption (Crabbe et al., 2011; Huynh et al., 2019; Rhodes et al., 2005; Simms et al., 2008; Sprow and Thiele, 2012; Thiele and Navarro, 2014). Parallel advances in high-density electrophysiological recordings (e.g., Neuropixels), multiphoton microscopy, and widefield calcium imaging now expand interrogation of brain function in awake, behaving animals across the spatial scale of information processing relevant for alcohol-related behaviors (Siciliano and Tye, 2018). These techniques allow functional activity and structural measurements of subcellular compartments (e.g., axons and dendrites), single cells (neuronal and non-neuronal glia cell bodies), and the network ensemble activity of hundreds to thousands of neurons simultaneously across multiple brain areas. Moreover, they open the possibility of longitudinally tracking the microscopic activity of single cells and macroscopic activity of individual brain areas across days to weeks of ethanol consumption, allowing for the systematic analysis of how neuronal activity changes with repeated ethanol intake. While these techniques require head-fixation, paradigms for voluntary drinking in head-fixed mice have not been characterized.

Here, we present a paradigm for voluntary, head-fixed ethanol drinking (HFD) as a modified extension of the well-studied “drinking-in-the-dark” (DID) paradigm (Rhodes et al., 2005; Thiele et al., 2014; Thiele and Navarro, 2014). Mice transfer binge-like drinking behavior from the classic DID paradigm in their home cages to head-fixation, thereby allowing alcohol studies with cutting-edge neuroscientific techniques that require head fixation (also see (Timme et al., 2023)). As an example application of this paradigm, we recorded PFC population activity with single-cell resolution using two-photon calcium imaging during head-fixed ethanol consumption. Drinking had heterogenous effects on single neuron activity at slow and fast time scales but did not significantly affect pairwise neuronal correlations. Together with traditional freely-moving paradigms, this new complementary approach will provide additional insights into the cellular- and circuit-specific mechanisms that contribute to the development and treatment of AUD.

## 2. Materials and Methods

### 2.1. Animals

Behavioral experiments were performed on male and female C57/BL6J mice maintained on a reversed light/dark circadian cycle with *ad libitum* access to standard mouse chow and water. Mice of either sex were ∼10 weeks at the start of behavioral experiments (mean 9.84, range 6-21 weeks). For two-photon calcium imaging experiments, animals expressed GCaMP6f under the CaMKII promoter and were generated by crossing the commercially available Camk2a-Cre (005359, Jackson) and Ai148D (030328, Jackson) mouse lines, both on a C57/Bl6 background. Imaging studies were done on ∼P60 male mice. Behavioral experiments were performed at Rutgers University and imaging experiments were done at MIT. All animal procedures were performed in strict accordance with protocols approved by the MIT Division of Comparative Medicine and Rutgers Comparative Medicine Resources and conformed to NIH standards.

### 2.2. Surgical procedures – headplate implant

Surgeries were performed under isoflurane anesthesia (4% induction, 1-3% maintenance). Body temperature was maintained at 37.5°C via a heating pad integrated into the base of the stereotaxic frame and a temperature controller (53800, Stoelting). Mice were given a subcutaneous injection of extended-release buprenorphine (3.25mg/kg) before surgery to provide analgesia for up to 72 hours post-surgery; meloxicam (10 mg/kg) was provided if additional analgesia was required during the recovery period. Anesthetized mice were head-fixed in a stereotaxic frame (51500D, Stoelting). Scalp hair was removed using a depilatory cream (Nair) and the scalp was disinfected using alternating scrubs with betadine and 70% ethanol solution. A portion of the scalp was removed and conjunctive tissues cleared after treatment with 3% hydrogen peroxide. The skull was abraded with a dental drill. A custom-designed headplate (eMachineShop) was placed over the skull and adhered in place using dental acrylic (Metabond, Parkell) mixed with black ink. Animals were then allowed to recover in their own cage with a warm water blanket and moistened food chow. Mice were singly housed for the remainder of the experiment and recovered from surgery for at least one week before beginning experiments.

### 2.3. Surgical procedures – chronic window implantation

The procedure for chronic window implantation was similar to headplate implantation as described in section 2.2 but with a few modifications. Surgeries were performed under isoflurane anesthesia (3% induction, 1-3% maintenance) and body temperature was maintained at 37.5°C using a temperature controller (ATC2000, World Precision Instruments). Animals were dosed with slow-release buprenorphine (0.1mg/kg) prior to surgery, and meloxicam (1mg/kg) every 24 hours post-surgery for 72 hours or until fully recovered. After exposing the scalp, a 3mm diameter craniotomy was drilled centered over the left ACC/M2 region (from bregma, AP: 1.0mm, ML: 1.0mm). A chronic cranial window, consisting of a 5mm diameter coverslip glued to two 3mm coverslips (Warner Instruments) with optical UV-cured adhesive (61, Norland) was then implanted. The window was carefully lowered with the 5mm coverslip on top and firmly held in the craniotomy using the stereotax arm and adhered to the skull using dental acrylic mixed with black ink (Metabond, Parkell). Once the dental acrylic had cured around the cranial window, the headplate was implanted using dental acrylic as described in 2.2. 3 mice in the 1-cycle paradigm received intracranial injection of an AAV virus expressing CaMKII-ChR2 in the ACC and were implanted with a bilateral optic fiber cannula above the injection. Data is presented from sessions before any optogenetic manipulations.

### 2.4. Drinking in the dark (DID) paradigm

Following recovery from surgery, mice were acclimated to ethanol consumption using the drinking in the dark (DID) paradigm. Mice for behavioral experiments were given 1 or 2 cycles of DID. For each DID cycle, mice had access to ethanol for 2 hours on days 1-3 and for 4 hours on day 4; no ethanol was available on days 5-7. Mice were weighed daily, and their water bottles were replaced with 10mL drinking bottles fitted with a sipper tube (Amuza) containing 20% ethanol (v/v) diluted in the mouse drinking water 3 hours into their dark phase (ZT15). Tubes were weighed before being placed in each cage. A control cage without any mice was also fitted with an ethanol drinking bottle with a sipper tube to account for evaporation. Mice were then left alone for the duration of the DID period to consume the ethanol solution. Afterwards, ethanol drinking bottles were removed, weighed, and regular drinking water bottles returned. The amount of ethanol consumed was calculated by subtracting the final weight from the initial weight of each tube to get a session difference. The control tube difference was then subtracted from each mouse tube difference, and then consumption computed as ethanol consumed (g) per weight of the animal (kg). Mice for two-photon imaging experiments were similarly acclimated to ethanol consumption except they performed DID for 13 consecutive days with ethanol access for 3 hours on each day.

### 2.5. Head-fixed drinking paradigm

The rig for head-fixed drinking (HFD) experiments was custom designed using parts from Thorlabs. After DID exposure, mice for behavioral experiments were habituated to head-fixation in 30-minute sessions over 2 days. For the two-photon imaging experiments, mice were habituated to head-fixation during the last 5 days of DID drinking. Animals were head-fixed on an elevated platform with a lickspout delivering 20% ethanol (v/v) in mouse drinking water positioned within easy access for licking. The lickspout was made from a brass tube (3.97mm diameter, 8128, K&S Precision Metals) for two-photon experiments or a blunt 13-gauge needle (McMaster-Carr) for behavioral experiments. For lick detection, the brass tube was wrapped with conductive wire and connected to a capacitive sensor (P1374, Adafruit) integrated to a breadboard. Alternatively, the 13-gauge needle was connected via a wire to a transistor-based electronic circuit for lick detection (Huda et al., 2020; Slotnick, 2009). In either case, contact of tongue to the spout generated a voltage signal that was fed to an Arduino board (Uno Rev3, A000066, Arduino) as an analog input and recorded via custom MATLAB scripts. Ethanol delivery was initiated by custom MATLAB scripts that sent a digital signal via the Arduino to toggle a transistor on the breadboard (IRF520PBF, Digi-Key) and open a 12V solenoid (VAC-100 PSIG, Parker or 161K011, NResearch). Ethanol solution was maintained in a graduated syringe, gravity fed into the solenoid, and calibrated to deliver a small drop with each trigger (∼8µL for two-photon experiments; ∼5µL for behavioral experiments). Following each session, total ethanol delivered was compared between the graduated syringe and the calculated trigger volume to ensure accurate quantification of total ethanol delivered (for detailed example circuit diagram, see Fig. 1). In a subset of experiments, we placed a weigh boat underneath the spout to catch and weigh any spilled liquid (2-cycle paradigm). For these experiments, consumption values were adjusted for the spilled liquid.

**Figure 1.**
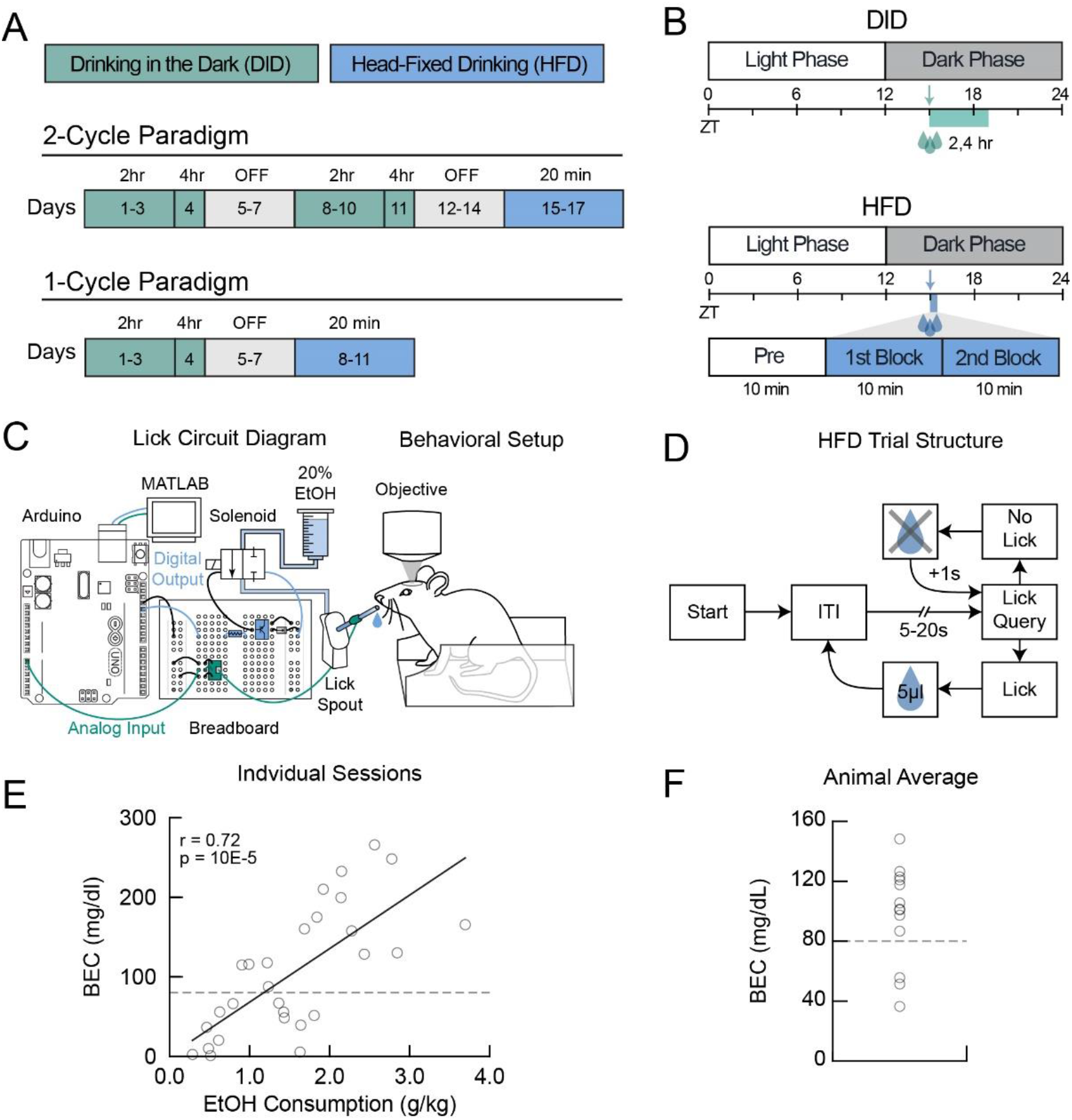
Paradigm for head-fixed ethanol consumption. **(A)** Mice first were acclimated to ethanol consumption using either 1 or 2 cycles of the ‘drinking in the dark’ (DID) paradigm with access to alcohol for 2 or 4 hours before undergoing head-fixed drinking (HFD) with access to alcohol for 20 minutes. **(B)** Mice started home cage drinking 3 hours into the dark phase of their circadian cycle. Head-fixed drinking started at the same time and consisted of three 10-minute blocks. A pre-drinking baseline block was followed by early and late drinking blocks. **(C)** Custom electronic circuit used for liquid delivery and lick detection during head-fixed drinking on a custom behavioral setup. **(D)** Trial structure for head-fixed drinking paradigm. **(E)** Relationship between head-fixed ethanol consumption and blood ethanol concentration quantified from plasma isolated from tail blood (2-4 sessions from each of 7 male and 6 female mice). **(F)** Session-averaged blood ethanol concentration for 13 mice.

Each HFD session consisted of three blocks: a 10-minute baseline pre-drinking period followed by two 10-minute drinking blocks. We head-fixed the animal and situated the spout in front of the mouth at the start of the session. The spout was aligned so it did not touch the animal but a protrusion of the tongue would be registered as a lick. Spout placement was inspected visually from multiple angles and via an infrared camera. Mice were free to lick the spout during the pre-drinking block, but licking did not deliver any liquid. Following the pre-drinking block, animals underwent two subsequent 10-minute drinking blocks in which licking the spout could deliver a drop of ethanol. We used a random interval schedule with a pseudorandom delay between drop deliveries (exponential distribution with a 10s mean and cut-offs at 5s and 20s). If the animal licked at least once since the prior delivery (or the beginning of the session for the first delivery), a drop of ethanol solution was dispensed. Otherwise, the delay was increased by 1s until a lick was made. Hence, the animal had to continually lick to trigger ethanol delivery. This design promoted licking of the spout in order to obtain more ethanol, thereby limiting unconsumed ethanol delivery. Moreover, the self-paced design allowed us to assess the animal’s level of engagement with ethanol by quantifying the number of drops that were triggered. An infrared camera and infrared illumination source were used to record pupil responses during the paradigm; these data are not included in this study and may be reported elsewhere.

### 2.6. Measurement of blood ethanol concentration

Blood ethanol concentration (BEC) for behavioral experiments was quantified with an Analox AM1 Analyzer using plasma separated from tail bloods. Mice were given additional HFD sessions at the end of the experiment for collecting tail bloods. For two-photon imaging experiments, BEC was quantified using a previously described assay (Prencipe et al., 1987). Immediately after the last drinking session, animals were rapidly anesthetized by isoflurane, decapitated, and whole trunk blood collected in tubes lined with EDTA (BD Microtainer, 365974, Becton, Dickinson and Company) and placed on ice. Whole blood was then centrifuged at 3000xg for 10 minutes at 4°C. Separated plasma was then aliquoted and immediately stored at −80°C for further analysis. Quantification of BEC was performed using a colorimetric assay as previously described. Ethanol standards and plasma samples were diluted in sample reagent (all reagents obtained from Millipore Sigma): 100mM KH_2_PO_4_ (P3786), 100mM K_2_HPO_4_ (P3786), 0.7mM 4-aminoantropyrine (A4382), 1.7mM chromotropic acid (27150), 50mg/L EDTA (E4884), and 50mL/L Triton X100 (X100). Working reagent was created by mixing alcohol oxidase from *Pichia* (5kU/L, A2404) and horseradish peroxidase (3kU/L, 77332) with sample reagent and mixed with samples on a 96-well plate. Following 30 minutes of incubation at room temperature, the samples and standards were read on a standard plate reader (iMark, Bio-Rad Laboratories) at 595nm. Samples and standards were run 6x in parallel and BEC calculated according to the standard curve.

### 2.7. Two-photon calcium imaging

GCaMP6f fluorescence from neuronal somas was imaged through a 16x/0.8 NA objective (Nikon) using resonance-galvo scanning with a Prairie Ultima IV two-photon microscopy system. Image frames were collected as 4-frame averages at 480x240 pixel resolution an acquisition rate of 16Hz. Excitation light at 900nm was provided by a tunable Ti:Sapphire laser (Mai-Tai eHP, Spectra-Physics) with ∼10-20 mW of power at sample. Emitted light was filtered using a dichroic mirror and collected with GaAsP photomultiplier tubes (Hamamatsu). Layer 2/3 GCaMP6f-expressing neurons were imaged with 1.5x optical zoom, 120-200μm below the brain surface. Neuronal activity in the anterior cingulate cortex (ACC) was collected at the following coordinates: ∼0.5mm AP, ∼0.5mm ML.

### 2.8. GaMP6f fluorescence signal processing

We used the software Suite2P (Pachitariu et al., 2017) for semi-automatic detection of neuronal somas from calcium imaging movies. Movies from the three imaging blocks were concatenated together and the non-rigid translation function in Suite2P was used to correct for x-y translations that may have occurred between blocks. Suite2P detects neuronal regions of interest (ROI) by clustering neighboring pixels with similar fluorescence time courses. Moreover, it provides for each detected neuron a neuropil mask which surrounds the detected ROI and excludes other detected neuronal ROIs. The automatically detected ROIs were manually curated using the GUI such that ROIs without clear structural evidence for neuronal somas were rejected and neurons missed by the algorithm were added manually.

To minimize the contribution of the neuropil signal to the somatic signal, corrected neuronal fluorescence at each time point t was estimated as F_t_ = F_soma_t_ – (0.3 x F_neuropil_t_) (Chen et al., 2013). The DFF (ΔF/F) for each neuron was calculated as ΔF/F(t) = F(t) – F_0_)/F_0_, where F_0_ represents the mode of the distribution of fluorescence values (estimated using the MATLAB function ‘ksdensity’). The resulting DFF trace was z-scored. We identified individual calcium events as transient increases in the z-scored DFF signal. Using the ‘findpeaks’ function in MATLAB, we detected events with minimum peak prominence of 2.5 z-scored DFF and minimum width of 3 imaging frames (∼200ms) at half-height of the event peak. All analyses either used the z-scored DFF or detected calcium event frequency and amplitude.

### 2.9. Analysis of change in neuronal activity with ethanol consumption

To determine how drinking affects ACC activity relative to before drinking, in later imaging sessions we introduced a 10 minute pre-drinking imaging block in which animals were allowed to lick the spout but no ethanol was delivered (note that the pre-drinking block was included for all behavioral experiments). We analyzed data from drinking sessions with imaging during both pre-drinking and drinking blocks. We had 13 such sessions from 4 male mice, which were included in the analysis of pre-drinking activity. 1 session had poor quality imaging data in the late drinking block and hence was excluded from the analysis.

We tested how ethanol consumption affects neuronal activity over the slow time scale of minutes across the entire imaging session. A challenge with this analysis is that the paradigm has no trial structure, making it difficult to align activity to specific events and statistically compare how drinking affects activity. While one strategy is to compare inter-event intervals between pre-drinking and drinking blocks, we reasoned that the sparse cortical activity observed in individual blocks would be a limiting factor and produce false negatives. Hence, we instead addressed this issue by devising a shuffle test. This test circularly shifted traces of detected calcium events in time by a random amount in intervals of 30s, thus maintaining the local temporal structure of activity while randomizing the time at which it occurred. We reiterated this process 1000 times for each neuron. On every iteration, we computed the difference in event frequency between pre-drinking and 1) the first drinking block; and 2) the second drinking block. This allowed us to generate null distributions for the difference in event frequency expected by chance given the overall activity level of the neuron. The two-tailed p-value for each drinking block was computed as the proportion of activity changes in the null distribution that were as or more extreme than the experimentally observed change on either side of the distribution. Neurons with p < 0.05 for either drinking block were considered significant and classified as drinking increased or drinking decreased.

We also tested how individual ACC neurons were modulated on a faster time scale. We aligned neuronal activity to the time of ethanol delivery and compared responses 1s before and after delivery using a one-tailed Wilcoxon signed-rank test to identify delivery increase and decrease neurons. Neurons with p < 0.01 were considered significant.

### 2.10. Pairwise neuronal correlation analysis

We assessed the effect of ethanol consumption on Pearson correlations between the z-scored DFF traces for all unique pairs of neurons in each recording session. In the drinking block animals lick to consume ethanol, but this licking is greatly reduced in the pre-drinking block. Hence, to facilitate comparison between drinking and pre-drinking blocks, activity 3s after ethanol delivery in the drinking blocks was not included in the analysis. Correlations were computed using the last 5 minutes of the pre and late drinking blocks. Correlations of all simultaneously recorded unique neuronal pairs were compared for the pre-drinking and drinking blocks. We also quantified pair-wise correlations as a function of the distance between neurons in each pair. We calculated the Euclidean distance between each pair of neurons as the length of a straight line connecting the center coordinates of each neuronal ROI. We defined proximal pairs as neurons less than 50 µm apart and distal pairs as neurons with more than 300 µm distance between them. We also analyzed whether the overall level of activity in the pair of neurons is related to ethanol’s effect on pair-wise correlations. We did a median-split for the average event frequency for each pair and compared pair-wise correlations separately for pairs with high and low levels of activity. Lastly, we quantified and compared the proportion of neuron pairs with significant positive or negative pair-wise correlations during pre-drinking and post-drinking (p < 0.05).

## 3. Results

### 3.1. A paradigm for ethanol consumption in head-fixed mice

We first acclimated mice to ethanol consumption (20% v/v in standard drinking water) in their home cages using the ‘drinking in the dark’ (DID) paradigm which promotes binge-like levels of ethanol consumption. Each cycle (week) of DID consisted of ethanol access for 2 hours on days 1-3 followed by 4-hour access on day 4 (**Fig. 1A,B**). After completion of DID drinking, mice were habituated to head-fixation for 2-3 days. We tested two experimental designs for assessing head-fixed drinking, with one group receiving 1 cycle of DID and the other group receiving 2 cycles of DID (**Fig. 1A**). For both groups, DID was followed by head-fixed drinking (HFD). This allowed us to assess how home cage drinking history affects head-fixed drinking. Moreover, in a subset of mice we assessed head-fixed consumption of 3% sucrose to compare drinking patterns with ethanol.

We custom designed software and hardware to assess ethanol consumption behavior in head-fixed mice. Liquid delivery was controlled via a solenoid valve, which was calibrated daily to deliver ∼5µL drops via a lick spout (**Fig. 1C**). The software controlled liquid delivery such that drops were dispensed with a pseudorandom delay between 5-20s with a mean delay of 10s (**Fig. 1D**). The system monitored the licking behavior in real-time to deliver drops only if the animal licked at least once since the previously delivered drop. Liquid delivery was delayed by 1s until the animal made at least one lick (**Fig. 1D**). This design minimized delivery of unconsumed drops of ethanol and allowed mice to self-pace their consumption.

Mice were head-fixed for 30 mins on each day of HFD sessions. We positioned the spout in close proximity to the mouth to facilitate licking and consumption. The first ten minutes constituted the pre-drinking block during which no ethanol was delivered (**Fig. 1B**). We included this block for future experiments to allow quantification of neuronal activity and/or physiological parameters in the absence of ethanol. Mice could consume ethanol during the subsequent 20-minute arbitrarily split into 1^st^ and 2^nd^ 10-minute blocks. We designed our HFD paradigm such that ethanol delivery is contingent on licking and consumption. Rarely, we observed spilled liquid from the spout. We investigated this issue in the 2-cycle cohort by weighing spillage of unconsumed liquid from the spout. The average spillage was minimal; we observed spillage in 3 sessions across 55 sessions from 15 mice. We quantified blood ethanol concentration (BEC) from tail blood collected from a subset of mice after completion of HFD sessions. There was a linear relationship between consumption on single HFD sessions and BEC (**Fig. 1E**). Session-averaged consumption for individual mice showed pharmacologically relevant BEC levels for most mice (**Fig. 1F**). Hence, head-fixed mice consume ethanol to binge-like levels.

### 3.2. Comparison of home cage and head-fixed ethanol consumption

Overall, mice in both the 2-cycle and 1-cycle cohorts consumed more ethanol during the 4-hr DID session compared to HFD (**Fig. 2A-D**). However, normalizing total consumption by duration of the session showed the opposite pattern, with a substantially higher drinking rate during HFD than during DID (**Fig. 2E, G**). Hence, mice consume more ethanol per unit of time during HFD. Previous work shows sex differences in DID home cage ethanol consumption, with female C57/BL6J mice consuming more ethanol than males (Radke et al., 2021). We found a similar sex difference in head-fixed drinking, with females drinking more than males in both cohorts (**Fig. 2F, H**). Comparing sex-specific consumption in the two HFD paradigms using a two-way ANOVA showed a significant main effect of sex (F(1,26) = 16.84, P = 4x10^-4^) but not for type of paradigm (F(1,26) = 2.43, P = 0.13). There was no interaction between sex and HFD paradigm (F(1,26) = 0.23, P = 0.64). Hence, mice consumed similar amounts of ethanol during HFD regardless of 1 or 2 weeks of DID ethanol exposure.

**Figure 2.**
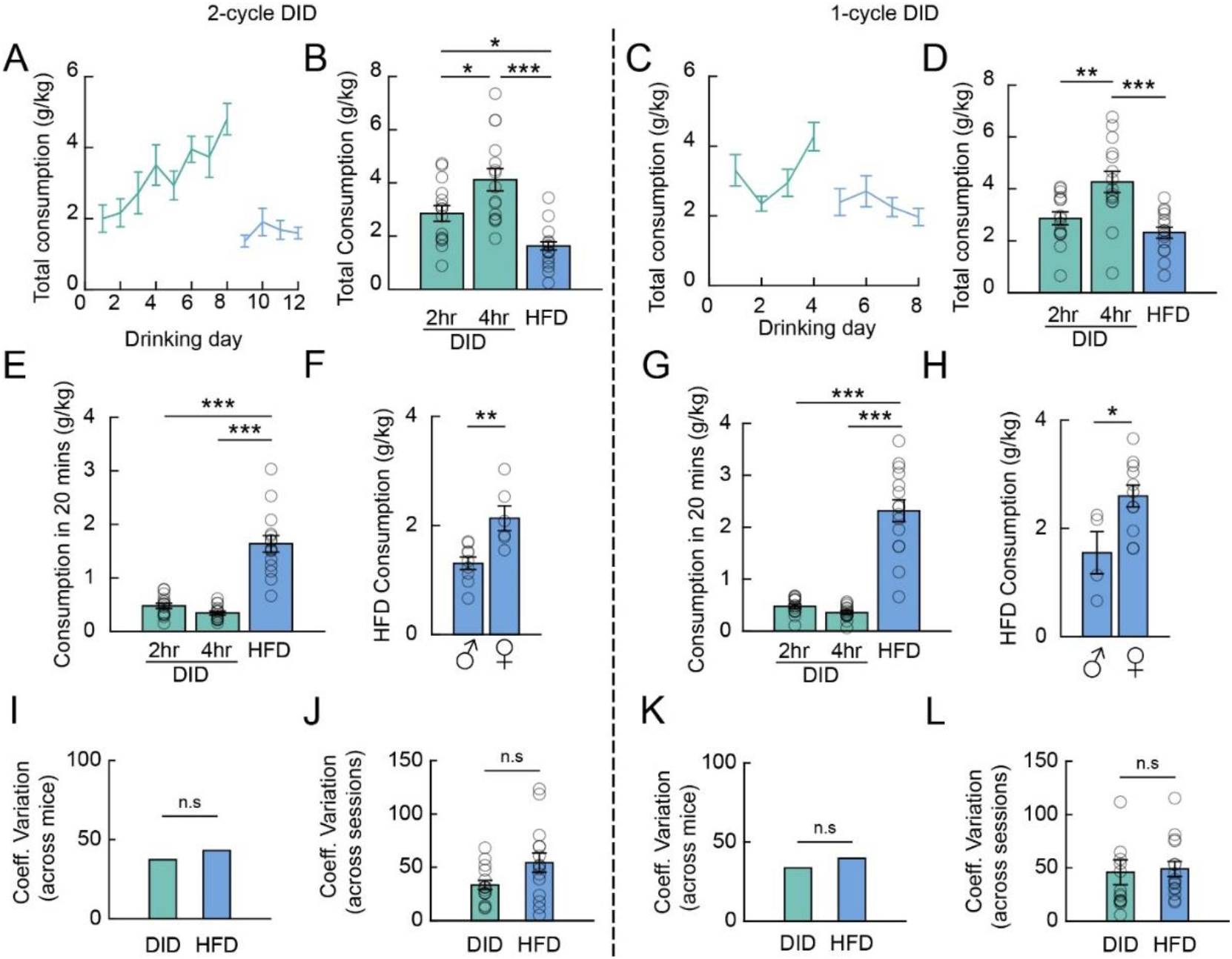
Comparison of home cage and head-fixed ethanol consumption. **(A)** Total ethanol consumption (g/kg) during home cage (green) and head-fixed (blue) drinking for mice in the 2-cycle paradigm. **(B)** Mean total consumption during 2-hr DID, 4-hr DID and 20 minutes of HFD (n=15 mice; one-way ANOVA main effect of paradigm type, F(2,42) = 16.15, P = 6.0e^-6^; Tukey HSD multiple comparisons, 2hr vs. 4hr: P = 0.016, 95% C.I. = [−2.33, −0.20]; 2hr vs. HFD: P = 0.021, 95% C.I. = [0.16, 2.29]; 4hr vs. HFD: P = 3e^-06^, 95% C.I. = [1.43, 3.55]. **(C)** Mean ethanol consumption during the 1-cycle paradigm. **(D)** Comparison of total consumption during 2-hr DID, 4-hr DID, and HFD drinking for 1-cycle paradigm (n = 15 mice; one-way ANOVA main effect of paradigm type, F(2,42) = 11.22, P = 0.0001; Tukey HSD multiple comparisons, 2hr vs. 4hr: P = 0.0053, 95% C.I. =[−2.44, −0.37]; 2hr vs. HFD: P = 0.41, 95% C.I. =[−0.49, 1.58]; 4hr vs. HFD: P = 0.0001, 95% C.I. = [0.92, 2.98]). **(E)** Data in **B** normalized to 20 minutes (n = 15 mice; one-way ANOVA main effect of paradigm type, F(2,42) = 55.75, P = 1.5e^-12^; Tukey HSD multiple comparisons, 2hr vs. 4hr: P = 0.59, 95% C.I. = [−0.19, 0.46]; 2hr vs. HFD, P = 2.4e^-10^, 95% C.I. = [−1.4877,-0.83311]; 4hr vs. HFD, P = 1.1e^-11^, 95% C.I. = [−1.62,-0.97]). **(F)** Head-fixed ethanol consumption in 2-cycle paradigm sorted by sex (n = 9 male and 6 female mice; unpaired t-test, t(13) = −3.5834, P = 0.003). **(G)** Data in **D** normalized to 20 minutes (n = 15 mice; one-way ANOVA main effect of paradigm type, F(2,42) = 75.8; P = 1.2e^-14^; Tukey HSD multiple comparisons, 2hr vs. 4hr: p = 0.78, 95% C.I. = [−0.31, 0.56]; 2hr vs. HFD: P = 1.4e^-12^, 95% C.I. =[−2.28, −1.41]; 4hr vs. HFD: P = 1.9e^-13^, 95% C.I. =[2.40, −1.53]). **(H)** Consumption in the 1-cycle paradigm sorted by sex (n = 4 male and 11 female mice; unpaired t-test, t(13) = −2.58, P = 0.023). **(I)** Across animal consumption variability. Coefficient of variation calculated across mice for mean consumption on 2-hour sessions of week 2 DID. For head-fixed drinking, it was calculated across mean head-fixed consumption over the first 3 days of HFD (Levene’s test, P = 0.19). **(J)** Across session consumption variability. Coefficient of variation calculated for each mouse across 2-hour sessions during week 2 of DID and across first 3 head-fixed drinking sessions (n = 15 mice; unpaired t-test, t(14) = −1.81, P = 0.092). **(K)** Same as I but for the 1-cycle paradigm (Levene’s test, P = 0.99). **(L)** Same as J but for 1-cycle paradigm (n = 15 mice; unpaired t-test, t(14) = −0.22, P = 0.83). Significance denoted as *p < 0.05; **p < 0.01; ***p < 0.005. All error bars are standard error of the mean.

We observed substantial across-animal and across-session consumption variability during HFD (**Supplementary Fig. 1A**). We compared variability during DID and HFD consumption to test if head-fixation uniquely contributes to consumption variability. Variability was calculated as the coefficient of variation using consumption values on 3 days of DID (2-hr sessions) and first 3 days of HFD. For both cohorts, the variability across mice was similar for average ethanol consumption during DID and HFD (**Fig. 2I, K).** Moreover, the mean across-session variability was also similar between DID and HFD (**Fig. 2J, L**). Therefore, consumption variability in the head-fixed paradigm likely represents biological variability in ethanol consumption behavior.

The addition of quinine to ethanol is often used to assess aversion-resistant drinking as a preclinical model of compulsive-like ethanol intake (Radke et al., 2021). We examined aversion-related modulation of head-fixed drinking in the 1-cycle group by adulterating the ethanol solution with increasing concentrations of quinine over 4 days after standard HFD (0.25 – 1mM in 0.25mM increments; **Supplementary Fig. 1B**). As expected, there was a significant decrease in HFD ethanol consumption with quinine adulteration (**Supplementary Fig. 1C**). These results suggest that the HFD paradigm is suitable for studying neural correlates of aversion-resistant drinking (also see Timme et al., 2023).

### 3.3. Frontloading behavior during head-fixed drinking

Our analyses thus far show similarities between DID and HFD for sex-dependent drinking and consumption variability. During home cage drinking, a subset of mice frontload and consume ethanol at a faster rate at the beginning of a drinking session (Ardinger et al., 2022). We analyzed the drinking pattern in HFD sessions to evaluate whether head-fixed mice also show front-loading behavior. Mice self-pace ethanol delivery in the HFD paradigm by licking the spout to trigger additional drops. We quantified the proportion of total drops that were dispensed in 2-min bins across 20 minutes of drinking (**Fig. 3A, B**). For both 2-cycle and 1-cycle groups, mice triggered more ethanol deliveries during the first 10 mins of the session compared to the last 10 minutes (**Fig. 3C**, D). We computed a front-loading score by normalizing the number of drops delivered in the first 10 mins by deliveries in the last 10 mins. There was no relationship between the frontloading score and the amount of ethanol consumed during HFD (**Fig. 3E, F**). This is consistent with observations during home cage drinking showing similar levels of total ethanol intake in frontloading and non-loading mice (Timme et al., 2023). These results show that head-fixed mice frontload ethanol.

**Figure 3.**
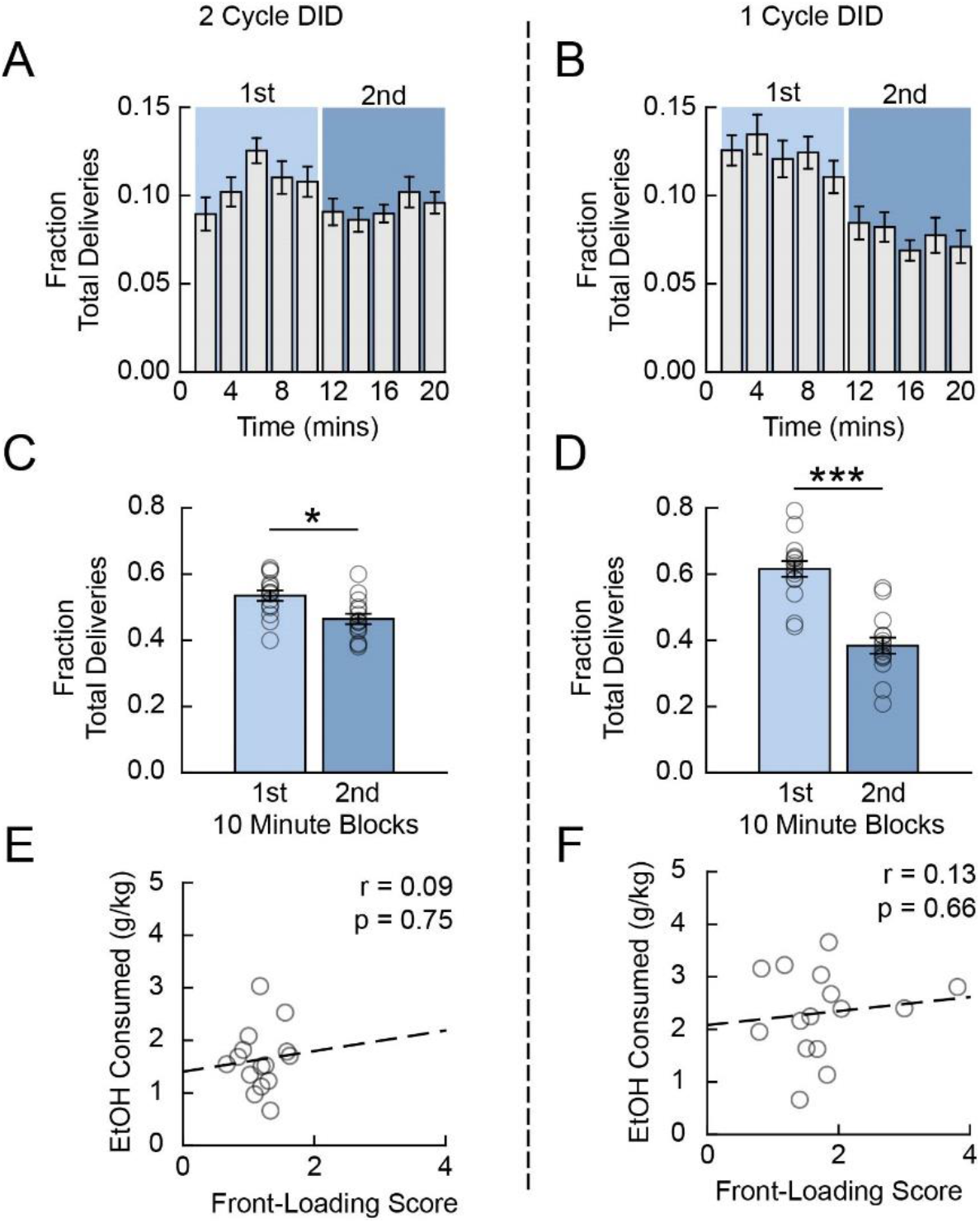
Frontloading behavior during head-fixed drinking. **(A, B)** Fraction of ethanol drops delivered in 2-minute bins across 20 minutes of head-fixed drinking in the 2-cycle (A) and 1-cycle (B) paradigms. **(C)** Comparison of ethanol consumption during early (1^st^ 10 minute) and late (2^nd^ 10 minute) drinking blocks (n = 15 mice; paired t-test, t(14) = 2.25, P = 0.041). **(D)** Same as (C) but for the 1-cycle paradigm (n = 15 mice; paired t-test, t(14) = 4.83, P = 2.7e^-4^). **(E, F)** Pearson’s correlation of front-loading score against head-fixed consumption for 2-cycle (E) and 1-cycle (F) DID paradigms. **(G)** Comparison of front-loading for mice in the two paradigms (n = 15 mice for each paradigm; two-sample t-test, t(28) = −2.75, P = 0.010. Significance denoted as *p < 0.05; **p < 0.01; ***p < 0.005. All error bars are standard error of the mean.

### 3.4. Licking dynamics during head-fixed ethanol consumption

Given the similar level of consumption observed in the two groups of HFD mice, we focused our remaining analysis on mice in the 1-cycle cohort. The HFD paradigm allows high temporal resolution tracking of licking behavior. We analyzed licking dynamics to better understand ethanol consumption behavior during head-fixation. Surprisingly, we observed licking during the pre-drinking period even when no ethanol was available for consumption, although there was significantly more licking during the drinking period (**Fig. 4A-C**). This pre-drinking licking is unlikely to reflect the motivation to consume ethanol as there was no relationship between the pre-drinking licking frequency and the number of ethanol drops delivered in the subsequent drinking session (**Fig. 4D**). Expectedly, the licking frequency during the drinking period was correlated with drop delivery (**Fig. 4E**), demonstrating that mice modulate their licking to acquire varying levels of ethanol during HFD.

**Figure 4.**
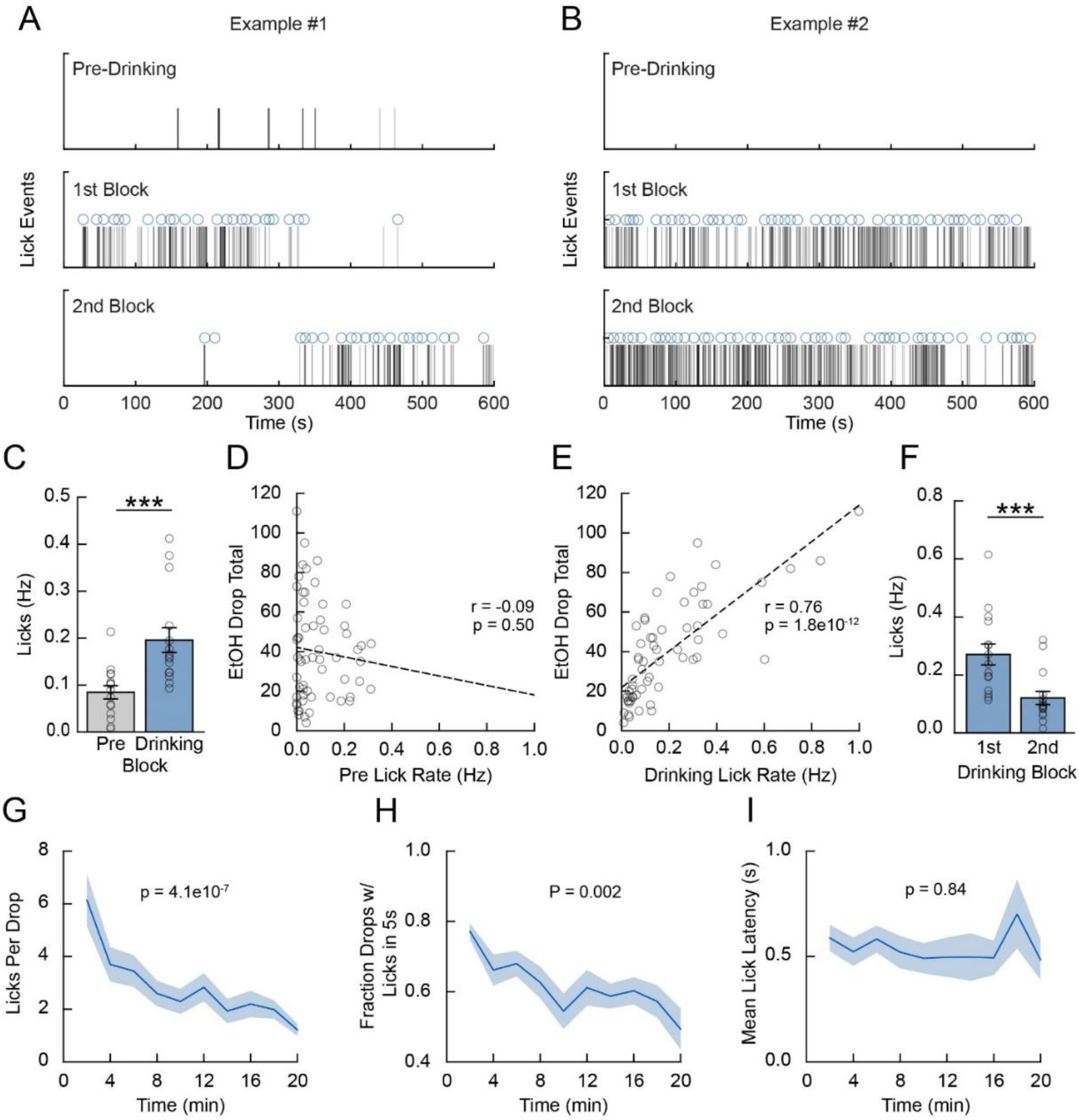
Licking dynamics during head-fixed ethanol consumption. **(A, B)** Licking activity on two example head-fixed drinking days. **(C)** Comparison of licking frequency during pre-drinking baseline period and during drinking (n = 15 mice; paired t-test, t(14) = −3.18, P = 0.007). **(D)** Correlation of pre-drinking licking frequency against total number of ethanol drops delivered for each HFD day (n = 60 sessions from 15 mice). **(E)** Correlation of drinking lick rate and number of delivered drops (n = 60 sessions from 15 mice). **(F)** Mean licking activity during 1^st^ and 2^nd^ 10-minute drinking sessions (n = 15 mice; paired t-test, t(14) = 5.02, P = 2.0e^-4^). **(G)** Mean number of licks in a 5s window for each ethanol delivery across 20 minutes of drinking in 2-minute intervals (n = 15 mice; one-way ANOVA main effect of time, F(9,140) = 6.02, P = 4.1e-07). **(H)** Fraction of delivered drops with mice licking within a 5s window (n = 15 mice; one-way ANOVA main effect of time, F(9,140) = 3.11, P = 0.002). **(I)** Mean latency to lick after ethanol drop delivery across drinking (n = 15 mice; one-way ANOVA main effect of time, F(9,139) = 0.54, P = 0.85). Significance denoted as *p < 0.05; **p < 0.01; ***p < 0.005. All error bars and shading are standard error of the mean.

Next, we analyzed how licking behavior evolves as a function of time during HFD sessions. The lick rate decreased in the second 10 mins of drinking compared to the first 10 mins (**Fig. 4F**). To investigate this further, we aligned licks to the time of individual ethanol deliveries and tracked across the session the number of licks elicited within a 5s window after each delivery. Average licks per drop decreased steadily over the course of the 20 mins drinking session (**Fig. 4G**). Inspection of individual licking traces showed that some drop deliveries were not immediately followed by licking (**Fig. 4A, B**). The fraction of deliveries with licks within a 5s window also decreased across the session (**Fig. 4H**). However, the mean latency to first lick after reward delivery did not change across the session (**Fig. 4I**). Taken together with the previous frontloading analyses (**Fig. 3**), these results show that mice trigger more drop deliveries and elicit more vigorous licking responses earlier in the drinking session.

### 3.5. Comparing head-fixed ethanol and sucrose consumption

We tested whether mice with on *ad libitum* water and food access (i.e., non-deprived) voluntarily consume sucrose during head-fixation. We reasoned that this would accomplish two goals: 1) it would allow us to compare head-fixed ethanol consumption to consumption of another rewarding liquid; and 2) it would allow future head-fixed studies to determine if neural determinants of consumption are specific for ethanol. Mice in the 1-cycle group were given a four-day break after their last ethanol/quinine head-fixed drinking session. Following this break, head-fixed mice self-administered 3% sucrose (w/v) during head-fixation with the same procedure used for ethanol consumption (**Fig. 5A**). We found that head-fixed mice readily consumed sucrose. Unlike for ethanol, sucrose consumption did not decrease across the 20-minute session and there was no difference in the number of delivered drops or the licking rate between the first and last 10 mins of drinking (**Figs. 3A-D**, **4F, 4G**, and **5B-D**). Although the number of licks per delivered drop of sucrose appeared to decrease across the drinking session (**Fig. 5E)** similar to ethanol (**Fig. 4G**), this effect was not statistically significant. Moreover, in contrast to ethanol (**Fig. 4H**), the proportion of drops with licking within 5s after delivery was similar across the session (**Fig. 5F)**.

**Figure 5.**
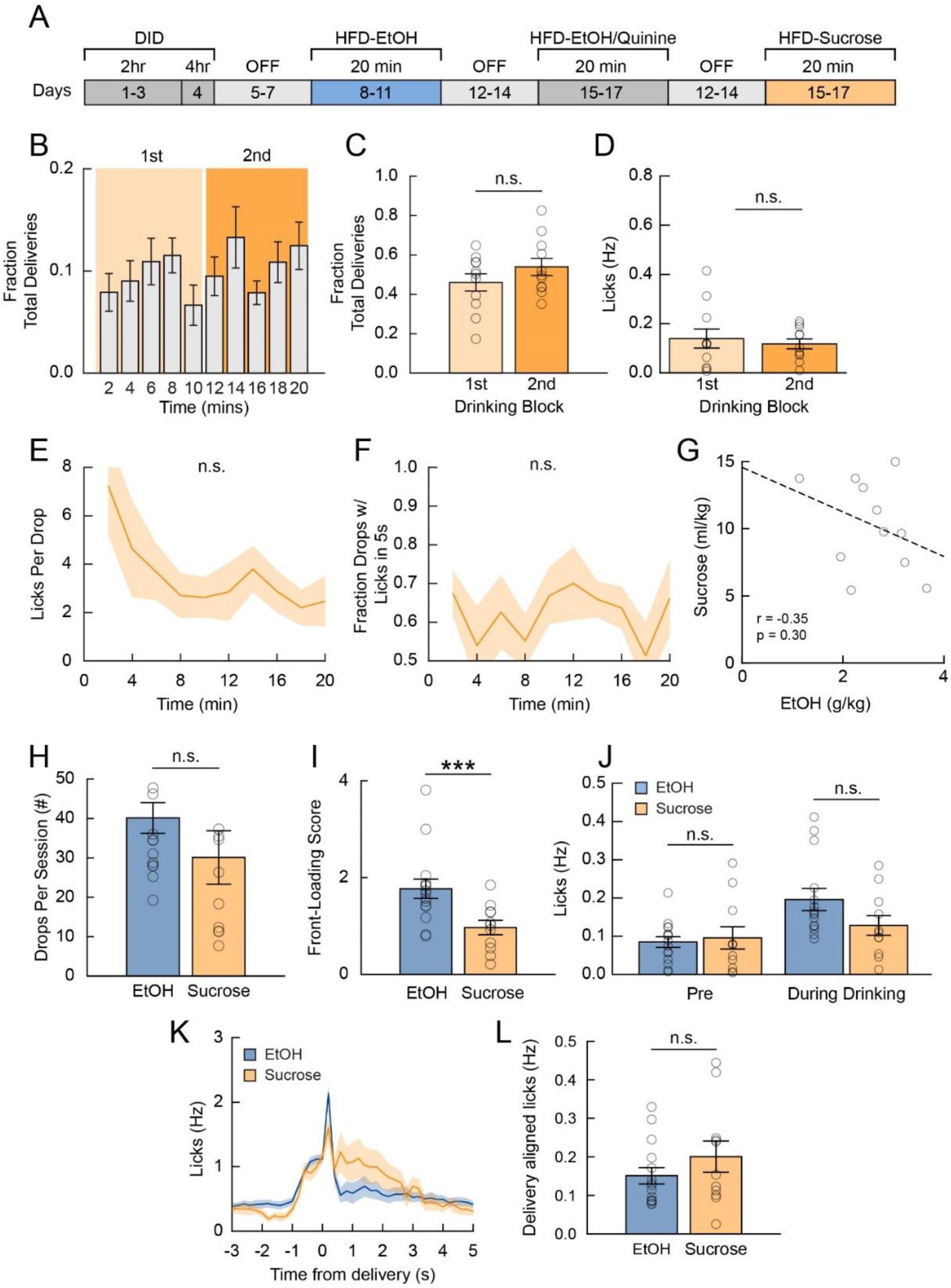
Comparison of ethanol and sucrose consumption during head-fixed drinking. **(A)** Timeline for experiments. **(B)** Fraction of deliveries across time during head-fixed sucrose consumption. **(C)** Comparison of sucrose deliveries triggered in 1^st^ and 2^nd^ sucrose drinking sessions. (n = 11 mice; paired t-test, t(10) = −0.91, P = 0.38). **(D)** Comparison of lick rates in 1^st^ and 2^nd^ sucrose drinking sessions (n = 11 mice; paired t-test, t(10) = 0.62, P = 0.55). **(E)** Mean number of licks in a 5s window for each sucrose delivery across 20 minutes of drinking in 2-minute intervals (n = 11 mice; one-way ANOVA main effect of time, F(9,87) = 1.33, P = 0.23). **(F)** Fraction of sucrose deliveries with a lick within 5s after delivery (n = 11 mice; one-way ANOVA main effect of time, F(9,87) = 0.64, P = 0.76). **(G)** Correlation of head-fixed ethanol and sucrose consumption. **(H)** Comparison of ethanol and sucrose drops delivered during head-fixed consumption (n = 15 and 11 mice for ethanol and sucrose, respectively; two-sample t-test, t(24) = 1.36, P = 0.19). **(I)** Front-loading score for ethanol and sucrose consumption (n = 15 and 11 mice for ethanol and sucrose, respectively; t(24) = 3.01, P = 0.006). **(J)** Comparison between ethanol and sucrose for pre-drinking and drinking licking rates (n = 15 and 11 mice for ethanol and sucrose, respectively; two-way ANOVA main effect of drink type, F(1,48) = 1.38, P = 0.25; main effect of session type, F(1,48) = 8.99, P = 0.0043; interaction between drink and session type, F(1,48) = 2.67, P = 0.11’ multiple comparisons with Tukey HSD, ethanol-pre vs. sucrose-pre: 95% C.I. = [−0.08 0.10], P = 0.99; ethanol-drinking vs. sucrose drinking: 95% C.I. = [−0.16 0.02], P = 0.21. **(K)** Licking dynamics for ethanol and sucrose head-fixed consumption. **(L)** Comparison of mean licking rate in a 3s window after sucrose and ethanol delivery (n = 15 and 11 for ethanol and sucrose; two-sample t-test, t(24) = −1.17, P = 0.25).

We directly compared ethanol and sucrose HFD data to further evaluate differences and similarities in the consumption of these liquids. Across mice, there was no relationship between the amount of ethanol and sucrose consumption (**Fig. 5G**). On average, mice delivered the same number of ethanol and sucrose drops (**Fig. 5H**) but showed less frontloading for sucrose than for ethanol (**Fig. 5I**). Comparison of licking behavior showed similar licking rates in the pre-drinking and drinking session for ethanol and sucrose (**Fig. 5J)**. Examining licking behavior aligned to the time of ethanol or sucrose delivery showed similar licking dynamics and rates for the two liquids (**Fig. 5K, L**). Together these analyses suggest that consumption and licking correlates of frontloading are more pronounced for ethanol than sucrose HFD.

### 3.6. Two-photon calcium imaging of the anterior cingulate cortex during head-fixed ethanol drinking

The prefrontal cortex is a key brain structure for high-level cognitive functions like attention, decision-making, and sensorimotor control. Acute ethanol intoxication is associated with deficits in these same functions, suggesting that drinking may affect prefrontal cortical activity. Previous *ex vivo* electrophysiological studies show that ethanol modulates a multitude of intrinsic electrophysiological and synaptic properties of neurons (Harrison et al., 2017; McCool, 2011). However, there is a paucity of data on how acute ethanol consumption affects the *in vivo* activity of prefrontal cortical networks during voluntary consumption. Our predominant goal in establishing a head-fixed ethanol consumption paradigm is to enable real-time interrogation of neural circuit function across acute and repeated ethanol consumption and during other ethanol-related behaviors. As an example application of our paradigm, we combined head-fixed drinking with two-photon calcium imaging to determine how ethanol consumption affects the activity of single prefrontal cortical neurons and network level interactions between neurons. Importantly, head-fixed mice consumed ethanol during two-photon imaging, achieving binge-like BEC levels (199.3±62.19 mg/dL, n=4 mice).

### 3.7. Effect of acute ethanol consumption on ACC single neuron activity

We used a transgenic mouse line (CaMKII-Cre x Ai148D on a C57/BL6 background) that targets GCaMP6f expression to excitatory pyramidal neurons in the cortex. Mice were implanted with a chronic window over the midline in the frontal cortex. We targeted recordings in the anterior cingulate cortex (ACC), a subdivision of the mouse prefrontal cortex that is accessible for two-photon imaging without brain lesioning optical implants (**Fig. 6A-C**). Although we could longitudinally track individual neurons across days, here we prioritized studying the effect of acute ethanol on large neuronal populations and analyzed the activity of >1400 recorded cells (n = 1471 neurons). Calcium events were detected from GCaMP6f fluorescence traces to quantify event frequency and amplitude in individual neurons. Inspecting the activity of individual neurons during pre-drinking and drinking showed heterogeneous effects of ethanol consumption on activity. The example neurons shown in **Fig. 6D** illustrate 3 types of observed responses over the time course of drinking: an increase, decrease, or no change in neuronal activity.

**Figure 6.**
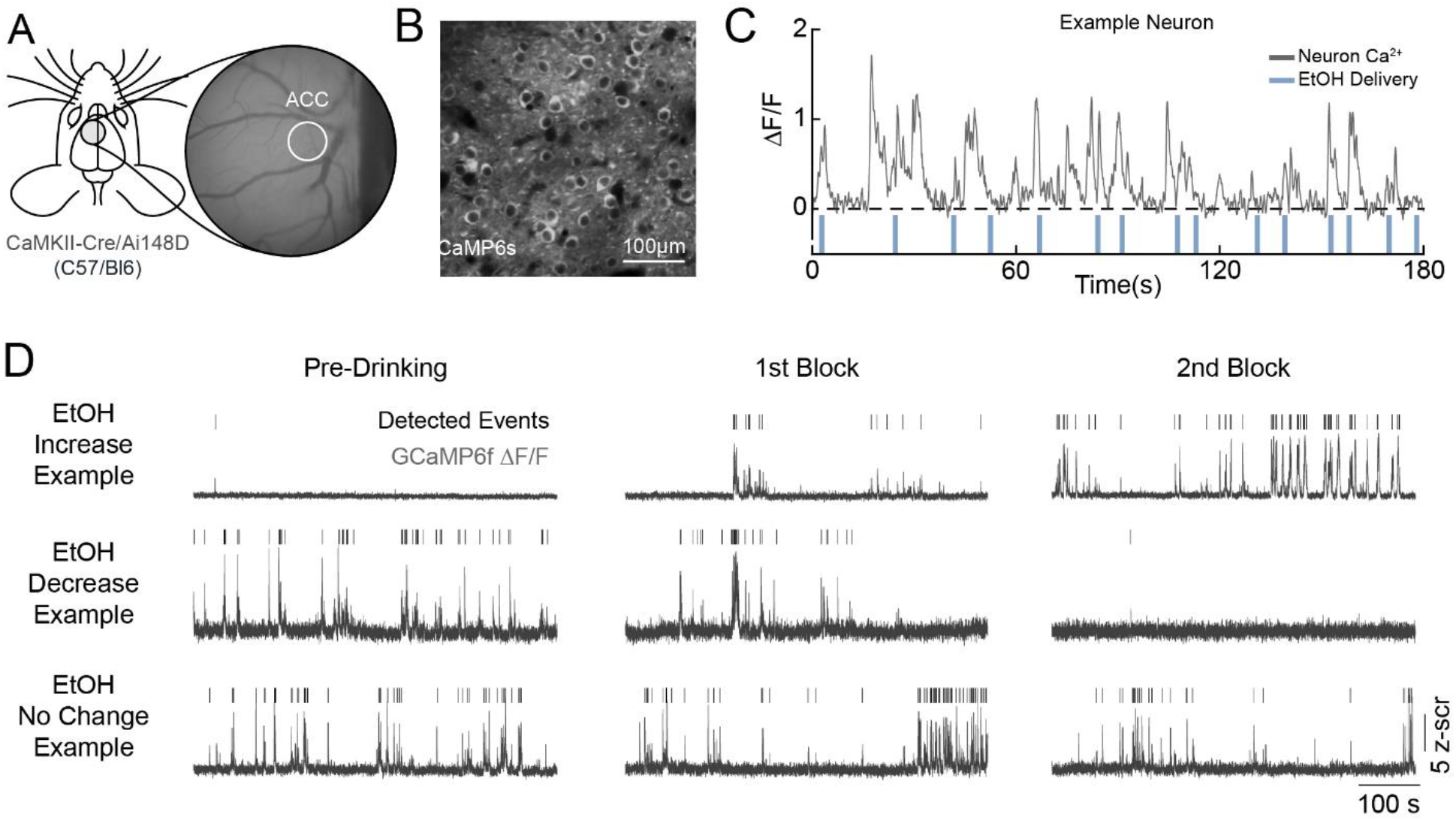
Two-photon calcium imaging of anterior cingulate cortex (ACC) activity during head-fixed ethanol consumption. **(A)** Chronic window implant site in the frontal cortex of transgenic mice expressing GCaMP6f in excitatory pyramidal neurons. **(B)** GCaMP-expressing ACC neurons. **(C)** Activity of an example neuron along with ethanol delivery times. **(D)** Three example neurons (rows) showing increase, decrease, or no change in activity from pre-drinking to the later drinking sessions.

To assess how the whole ACC population responds to alcohol consumption, we compared event rates of all recorded neurons during pre-drinking and the last 10 minutes of drinking. Overall, event rates were largely similar across these conditions (**Fig. 7A**). This lack of a population level effect is likely driven by heterogeneity in the response profiles of individual neurons, as suggested by the single neuron examples (**Fig. 6D**). The lack of a trial structure in this paradigm makes it difficult to statistically assess changes in activity levels for individual neurons in an unbiased way. We addressed this issue by performing for each neuron a shuffle test which compared the observed difference in activity during drinking and pre-drinking to the difference expected by chance given the overall activity in the entire recording period (see Methods). We found that alcohol increased and decreased the activity of a similar proportion of neurons **(Fig. 7B-E**); the activity of 11.6±1.4% of neurons was increased and the activity of 13.4±1.8% of neurons was decreased (p = 0.54, Wilcoxon signed-rank test; n = 12 sessions from 4 mice). Hence, despite ethanol’s effect on many intrinsic neuronal properties and on synaptic transmission, *in vivo* activity rates for the majority of ACC neurons are remarkably resilient against large changes with ethanol consumption.

**Figure 7.**
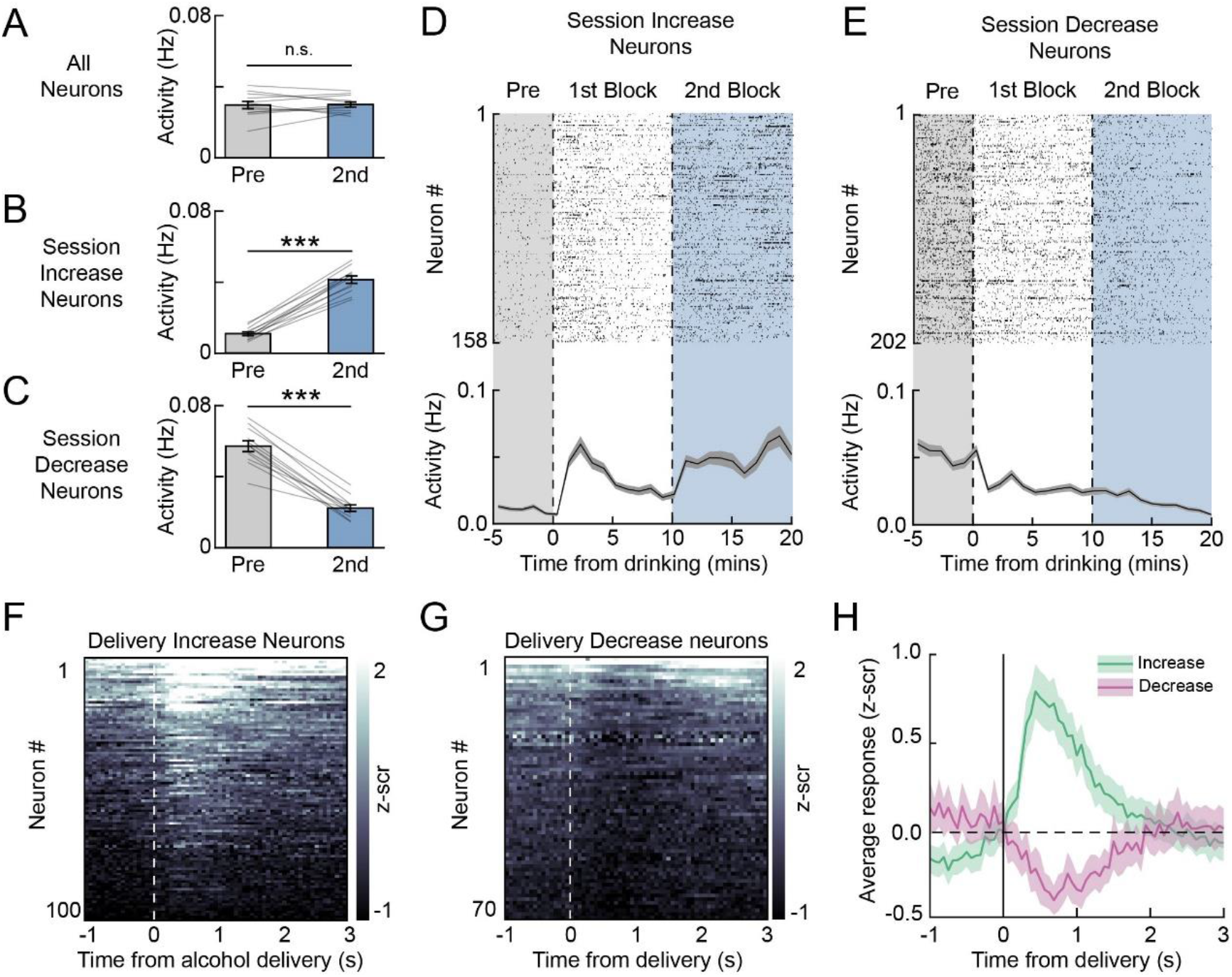
Effect of ethanol consumption on single neuron ACC activity. **(A)** DFF event rates during pre-drinking (gray) and drinking during 2^nd^ block (blue) for all neurons (P = 0.88, z = −0.16; n = 12 sessions from 4 mice) **(B)** Same as **A**, except for neurons with increased session activity (P = 0.002; z = −3.06; n = 12 sessions from 4 mice). **(C)** Same as **A,** except for neurons with decreased activity (P = 0.002; z = 3.06; n = 12 sessions from 4 mice). **(D)** Raster plot showing detected events for neurons with significantly increased activity across the drinking session (top). Rows show activity for individual neurons. Mean event rates in 1 min bins averaged across all increased activity neurons are shown below (shading is SEM). **(E)** Same as **D**, except for decreased activity neurons. **(F)** Color plot of the trial-averaged activity of neurons (rows) showing increased activity around the time of ethanol delivery. **(G)** Same as G, except for neurons showing decreased activity around ethanol delivery. **(H)** Average z-scored DFF across neurons with increased (green) and decreased (magenta) activity that are shown in (F) and (G), respectively.

The above analysis considers activity changes during drinking on the timescale of minutes, possibly reflecting the cumulative effect of ethanol consumption. Next, we aligned neuronal responses to individual ethanol deliveries and identified modulated neurons by comparing responses before and after delivery. Similar proportions of neurons had increased and decreased activity (6.6±3.8% & 4.3±3.5%, p > 0.05; **Fig. 2F-H**). These results show that in a small subset of ACC neurons, phasic responses are associated with consumption-related processes.

### 3.8. Effect of ethanol consumption on pairwise inter-neuronal correlations

The above analyses show that acute ethanol consumption modulates the activity of a subset of ACC excitatory neurons over slow (minutes) and fast (seconds) time scales. Information processing in cortical networks is shaped by correlated activity between neurons in addition to single neuron activity levels (Cohen and Kohn, 2011; Kohn et al., 2016). We took advantage of the large number of simultaneously recorded neurons in our dataset to test the effect of ethanol consumption on inter-neuronal correlations (**Fig. 8A**). For each recording session, we computed the Pearson correlation coefficient between the activity of unique pairs of neurons (**Fig. 8B**). Visualizing the activity of single example pairs showed diverse changes, with correlations increasing, decreasing, or being unaffected during the end of the drinking session (last 5 minutes of the second drinking block) compared to pre drinking (**Fig. 8B**). To examine this process at the population level, we compared correlations averaged over all unique pairs of neurons recorded simultaneously in single behavioral sessions. Overall, alcohol consumption did not significantly affect pairwise correlations (**Fig. 8C**).

**Figure 8.**
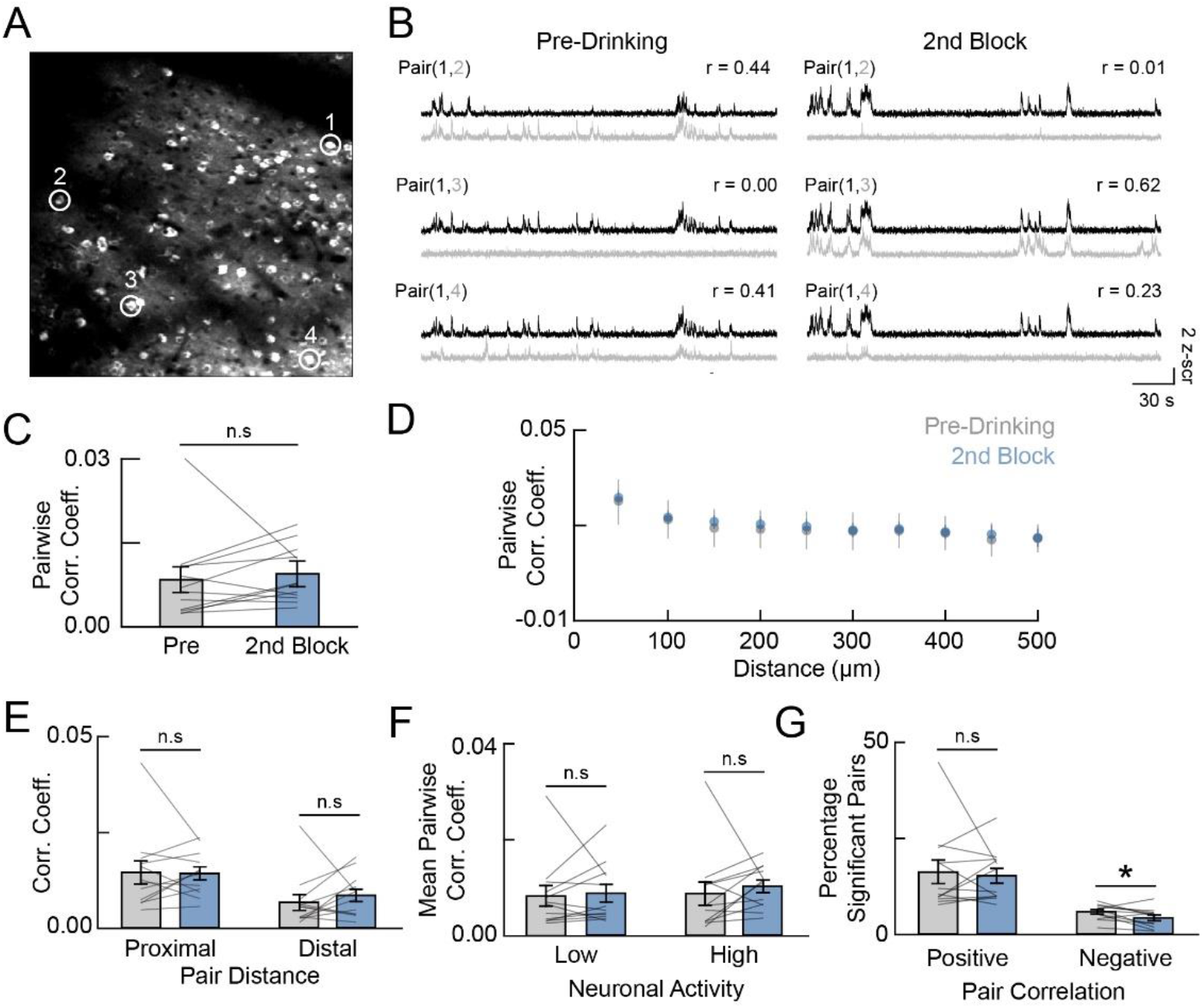
Effect of ethanol consumption on pairwise activity correlations between neurons. **(A)** Example field of view noting the spatial location of four representative neurons (white circles/numbers). **(B)** Representative activity of pairs of neurons shown in (A) for pre-drinking and the 2nd drinking blocks. **(C)** Mean pairwise correlation coefficients averaged for all unique neuronal pairs during pre-drinking and 2^nd^ drinking blocks (P = 0.18, z = −1.33, n = 12 sessions from 4 mice; Wilcoxon signed-rank test). **(D)** Pairwise correlation coefficients binned by the distance between neurons in each pair (bin size, 50µm). **(E)** Activity correlations of proximal (<50µm apart) and distal (>300µm apart) pairs during pre-drinking (gray) and 2^nd^ drinking (blue) blocks (proximal, P = 0.58, z = −0.55; distal, P = 0.21, z = −1.26). **(F)** Comparison of correlation coefficients of pairs with low and high activity, defined based on the median split of the calcium event frequency averaged across the neurons in a pair (low activity, P = 0.88, z = −0.15; high activity, P = 0.07, z = −1.80). **(G)** Percentage of neuronal pairs with significant positive or negative pairwise correlations (positive, P = 0.94, z = −0.08; negative, P = 0.041, z = 2.04). Error bars are SEM (n = 12 sessions from 4 mice). Significant testing with Wilcoxon rank-sum test; *p < 0.05; n.s., not significant.

Collapsing pairwise correlations across the whole recorded population may obscure subtle effects of ethanol on network-level interactions. We explored this possibility by breaking down pairwise correlations along several factors. Nearby cortical neurons share similar synaptic inputs, leading to a distance-dependence in pairwise correlations. In agreement, pairwise correlations generally decreased as a function of the distance between neurons in a pair, but there was no difference in this relationship with drinking (**Fig. 8D, E**). Neurons with high levels of activity may show higher pairwise correlations just by chance. Accounting for the average level of activity between neuron pairs did not show any differences (**Fig. 8F**). Lastly, we restricted our analysis only to neuronal pairs showing statistically significant activity correlations. The proportion of pairs with significant positive pairwise correlations was similar between pre-drinking and end of drinking. However, there was a small reduction in the proportion of neurons with negative pairwise correlations (**Fig. 8G**). Together, these analyses show pairwise correlations are largely unaffected by ethanol consumption.

## 4. Discussion

Neuroscientific research has benefitted from the rapid development of technologies to measure and manipulate neuronal and non-neuronal circuits. The rise of multiphoton imaging, high-density electrophysiological electrodes, fluorescence-based voltage sensors, and variety of other techniques have provided new insights into the spatiotemporal dynamics of neuronal computations important for behavioral control (Ji et al., 2016; Liu et al., 2022; Steinmetz et al., 2018; Tian et al., 2023). However, many of these tools currently require immobilization of the animal’s head. This methodological limitation necessitates the adaptation of current freely moving behavioral paradigms to head-fixed counterparts.

Here, we demonstrate the feasibility of a ‘head-fixed drinking’ (HFD) variant of the well-characterized ‘drinking-in-the-dark’ (DID) model of binge-like ethanol consumption. In our animal cohorts, mice consumed ∼1.5-2 g/kg of 20% ethanol over the course of 20 minutes, which represents an average rate of ∼0.1 g/kg/min. Comparatively, mice undergoing the traditional DID paradigm consume ∼3-4 g/kg over 2 hours, which corresponds to a rate of ∼0.025 g/kg/min (Thiele and Navarro, 2014). Therefore, although the overall ethanol volume consumed in HFD is less than DID, the rate is higher, resulting in pharmacologically relevant BEC levels for studying binge-like consumption. Given the rapid rate of ethanol consumption during head-fixation, this paradigm is a potential model for high-intensity binge drinking (Patrick and Azar, 2018), although further work is required to explore this possibility.

Despite the difference in intake rate, we found several similarities between home cage and head-fixed drinking. First, female mice consumed more ethanol during HFD than males, which is consistent with home cage drinking (**Fig. 2F, H**). This suggests that factors driving sex-differences in ethanol consumption behavior remain in effect with head-fixation. Second, session-to-session and animal-to-animal variability was similar between home cage DID and HFD drinking (**Fig. 2I-L**). Lastly, our self-paced design, where mice can withhold licking to stop ethanol deliveries, allowed us to assess the temporal pattern of ethanol consumption. We observed frontloading behavior during HFD such that consumption was skewed towards the beginning of the HFD session (**Fig. 3**). Importantly, there was no frontloading for sucrose (**Fig. 5C, D**). Such frontloading behavior has also been observed in home cage drinking as well as during operant responding for alcohol (Ardinger et al., 2022).

Head-fixation benefits several other detailed measurements of animal behavior (Ozgur et al., 2023). For example, using high resolution cameras, licking kinetics and microstructures can be readily recorded. Additionally, facial movements, whisking, and pupil diameter can be quantified using open-source, machine learning algorithms such as DeepLabCut (Nath et al., 2019). Head-fixation also allows for more controlled sensory environments that range in complexity from precise auditory and visual stimulation such as used for operant responding to closed-loop virtual reality environments. Controlling the sensory environment reduces trial-to-trial variability in contextual learning, operant conditioning, and sensorimotor decision-making. These behaviors (and others) have all been applied in the alcohol research field, but not within the head-fixed domain. Therefore, adapting these behavioral paradigms to head-fixed drinking will serve as a useful complementary approach to the freely moving counterparts and will allow application of recording modalities like two-photon imaging for assessment of circuit mechanisms.

Though HFD provides several benefits for some quantitative measures, it also presents a few experimental limitations. First, HFD requires the surgical implantation of a head-plate for head-immobilization. Custom stainless-steel head-plates can be manufactured relatively cheaply from open-source designs and can be recovered and sterilized for reuse across animals. Multiple surgical procedures for affixing the headplate to the animal’s skull have been published with the most common method being the use of dental cement (Carlsen et al., 2022; Holtmaat et al., 2012; Manita et al., 2022). Though headplate attachment is a relatively simple procedure, it does require substantial recovery by the animal especially if combined with a cranial window implant.

Second, HFD requires a platform with head-fixation bars that allows the animal to be supported while the head is immobilized. Custom head-fixation rigs can be built using readily available parts from opto-mechanical suppliers (e.g., Thor Labs) (Huda et al., 2020; Lee et al., 2022; O’Connor et al., 2010; Ozgur et al., 2023). The precise configuration will depend on the experimental parameters and measurements and may include a rotary encoder for measuring animal locomotion on a running wheel, an infrared camera and light source for pupil/face measurements, and stimulus presentation equipment (Ozgur et al., 2023). At a minimum, a lick spout and lick detection circuit must be built to deliver ethanol and record lick rates (Williams et al., 2018). Here, we use an Arduino UNO microcontroller to interface with MATLAB on the main experimental computer. Thus, despite the need for construction of a head-fixation rig and ethanol delivery circuit, the overall cost for HFD is considerably lower than many commercial behavioral systems.

A third, obvious limitation is the fact that the animal’s head is immobilized, which introduces potential confounds relating to stress levels (Juczewski et al., 2020). There are several ways to reduce stress associated with head-fixation (Barkus et al., 2022). The first method, employed here, is to habituate animals to handling and head-fixation. Habituation can limit the effect of these stimuli on the animal’s stress state, but of course this requires consistent exposure to the same experimental rig and/or experimenter. The number of habituation sessions can range from several days to several weeks depending on how important it is to reduce head-fixed stress for the experimental question (Juczewski et al., 2020; Russo et al., 2021). Furthermore, providing a running wheel allows mice to move even when the head is stationary. This can be achieved using either a flat disk or a light sphere levitating on air. Additionally, stress can be reduced by providing a dark, sound-dampened enclosure that separates the animal from the experimenter and limits visual, auditory, and olfactory stimuli that may induce stress responses. In addition to preventative measures for mitigating stress, quantifying the degree to which head-fixation affects different animals may help control for stress as a source of behavioral and physiological variability. In addition to measuring pupil dynamics and locomotion, blood samples can be acquired to assay for circulating stress hormones. In this way, heterogeneity in drinking behavior and physiology can be mapped onto potential differences in stress response to head fixation.

A critical consideration for implementing HFD is the precise positioning of the lick spout. It is recommended that a high-resolution fixed camera be used to record the relative positioning of the lick spout with each individual mouse. This reduces the variability in drinking due to unfamiliar lick spout placement. In addition, ethanol spillage, that is drops of alcohol triggered by licking but not consumed, can be a factor in drinking heterogeneity. The use of a high-resolution camera can help identify drops being delivered but not consumed. We placed a weigh boat underneath the spout to quantify unconsumed ethanol and found that spillage does occur but relatively infrequently. This also helps calibrate the solenoid delivery of ethanol by using both a volume release (measured by the graduated syringe containing the ethanol reservoir), the weight of the ethanol that lands on the weigh boat, the duration of time the solenoid was open (to deliver a single drop), and the total number of drops. A calibration table should be generated by varying the number of drops and the time the solenoid is open and interpolating a curve to the total delivery. Importantly, the BEC should be plotted against the measured consumption at least in a subset of animals or sessions in order to infer the relationship between consumption and BEC.

Here, we demonstrate that one experimental advantage of HFD is to apply two-photon microscopy to studying the heterogeneity of neuronal activity during epochs of alcohol consumption. We find subsets of excitatory neurons in layer 2/3 of the ACC that show either increased or decreased activity across ethanol consumption. Surprisingly, the population proportions of these subsets are relatively similar and the population average firing rate does not significantly change with ethanol consumption. This observation corroborates previous work in the rat prefrontal cortex using *in vivo* electrophysiology demonstrating significant heterogeneity of neuronal excitability in response to ethanol exposure using *in vivo* electrophysiology but no average change in firing rate at the population level (Linsenbardt and Lapish, 2015; Morningstar et al., 2020). We also found similar heterogeneity in acute neuronal responses, both positive and negative, to the delivery of alcohol, which may also represent licking activity, ethanol valence, behavioral and autonomic arousal, and/or other consumption-related processes. Further work is needed to map the heterogeneity of these responses to animal consummatory behavior including measures of sympathetic outflow.

The individual heterogeneity of ACC neuron activity in response to HFD could represent a broader change in population dynamics. Though we did not see an overall effect of ethanol consumption on firing rates, the correlated activity across the population could potentially be affected. However, by comparing the pair-wise correlations between neuron pairs, we did not find any significant change even when considering the spatial location of neurons and their overall activity levels. This suggests that in addition to having no net effect on firing rates, ethanol consumption does not broadly affect neuronal correlations in layer 2/3 of the mouse ACC. These mice only had a limited exposure history and it is possible that repeated ethanol consumption may more severely affect activity levels and correlations. Alternatively, these findings may point to systems-level mechanisms that maintain neuronal activity and correlations within a narrow setpoint and confer resiliency against large changes in activity levels.

A benefit of HFD and two-photon microscopy is the ability to longitudinally track individual neurons across days or weeks. Here, we analyzed single sessions across animals. Future experiments are needed to determine whether the individual firing rates and population correlations are stable over multiple sessions within the same animal (e.g., do neurons with increased firing rates in response to HFD show the same effect across days?). By tracking ethanol consumption and neuronal physiology longitudinally, cortical activity signatures may predict variable ethanol consumption within animals.

Another benefit of HFD and two-photon microscopy is the ability to quantify activity in genetically identified (e.g., cell-type specific expression of calcium indicators) or anatomically identified (e.g., anterograde/retrograde traces) subpopulations of neurons. We limited our analysis to CaMKII-expressing cells, which are predominantly excitatory in the cortex. However, increasing evidence suggests that inhibitory neuronal subtypes play important roles in the development of drinking behavior in the cortex (Dao et al., 2021; Fish and Joffe, 2022; Patton et al., 2023). By using dual-color calcium indicators, both excitatory and inhibitory populations can be simultaneously imaged using two-photon and HFD. Future work is needed to track the temporal evolution of these populations with the acquisition and maintenance of ethanol drinking behavior. Overall, we believe the HFD paradigm complements freely moving paradigms, allowing alcohol studies using activity recording and manipulation techniques not readily available in other models.

## Competing interest

The authors declare no conflicts of interest associated with this manuscript.

## Acknowledgements

This work was supported by grants from the National Institute of Mental Health (R00MH112855 to R.H.), Brain and Behavioral Research Foundation (R.H.), and National Institute on Alcohol Abuse and Alcoholism (K99AA028579 to G.O.S. and U01-AA025481 to E.M.V).

## CRediT authorship contribution statement

**Anagha Kalelkar:** Investigation, Software, Validation. **Grayson Sipe:** Conceptualization, Methodology, Investigation, Writing – original draft, Writing – review & editing, Visualization, Funding acquisition. **Ana Raquel Castro E Costa:** Investigation. **Ilka M. Lorenzo:** Investigation. **My Nguyen:** Investigation. **Ivan Linares-Garcia:** Software, Formal analysis. **Elena M. Vazey:** Methodology, Funding Acquisition. **Rafiq Huda:** Conceptualization, Methodology, Software, Formal analysis, Resources, Writing – original draft, Writing – review & editing, Visualization, Supervision, Project management, Funding acquisition.

## Data and code availability

Data and analysis code will be made available by the corresponding author upon request.

## Supplementary Figures

**Supplementary Figure 1.**
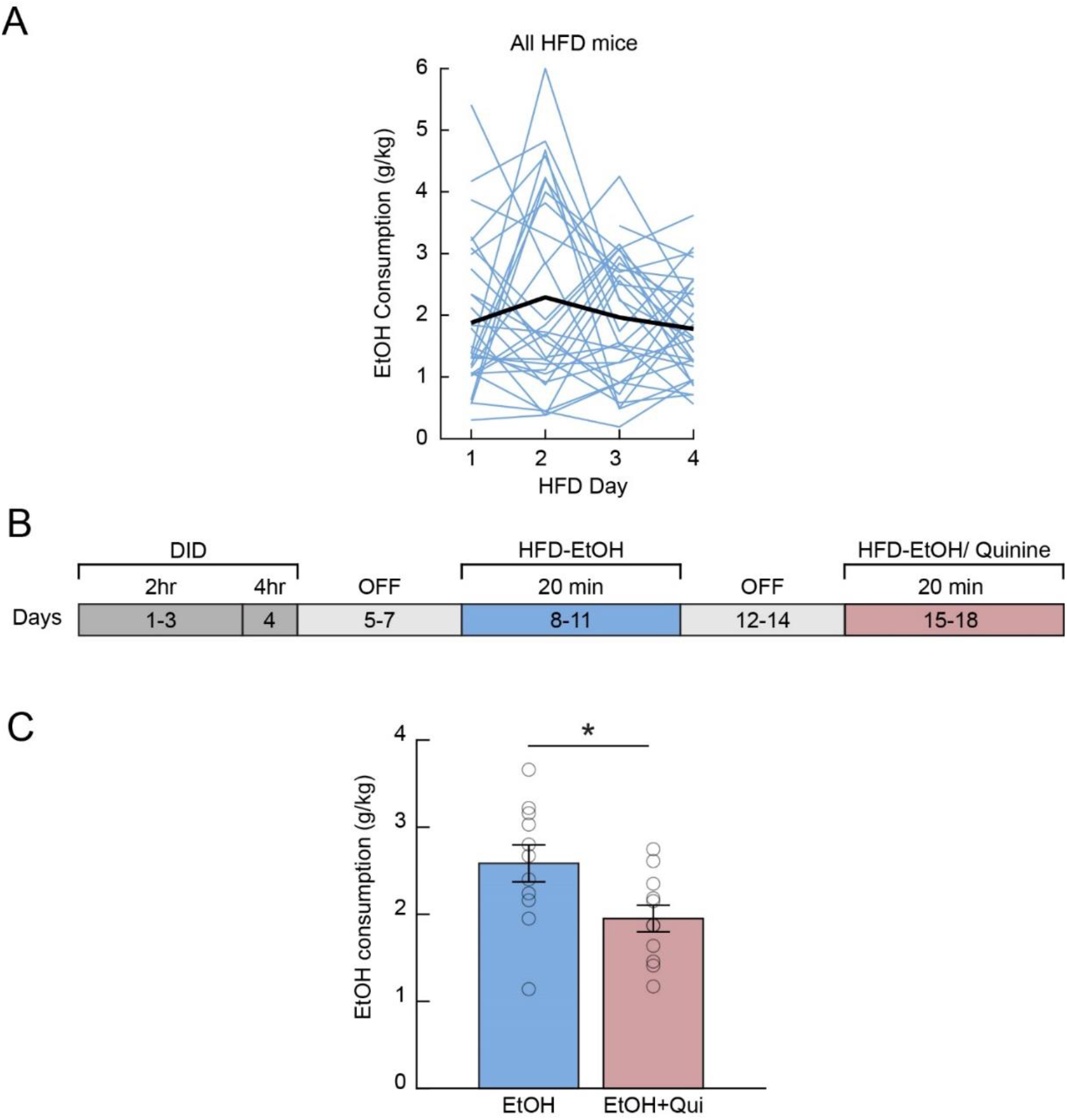
Within animal variability and the effect of quinine adulteration on HFD ethanol consumption. **(A)** Total ethanol consumption for individual animals across multiple days of HFD (n = 30 mice). **(B)** Experimental timeline for quine adulteration following standard HFD. **(C)** Ethanol consumption on day 11 of head-fixed drinking compared to days 16-18 when 0.5-1mM quinine was added to the 20% ethanol drinking solution. (n = 11 mice; paired t-test, t(10) = 2.52, P = 0.031).

## Notes

### Competing Interest Statement

The authors have declared no competing interest.

